# Insertions of codons encoding basic amino acids in H7 hemagglutinins of influenza A viruses occur by recombination with RNA at hotspots near snoRNA binding sites

**DOI:** 10.1101/2020.09.21.303073

**Authors:** Alexander P. Gultyaev, Monique I. Spronken, Mathis Funk, Ron A.M. Fouchier, Mathilde Richard

**Affiliations:** Department of Viroscience, Erasmus Medical Center, P.O. Box 2040, 3000 CA Rotterdam, the Netherlands; Group Imaging and Bioinformatics, Leiden Institute of Advanced Computer Science (LIACS), Leiden University, P.O. Box 9512, 2300 RA Leiden, the Netherlands

**Keywords:** Influenza virus, avian influenza, snoRNA, RNA recombination

## Abstract

The presence of multiple basic amino acids in the protease cleavage site of the hemagglutinin (HA) protein is the main molecular determinant of virulence of highly pathogenic avian influenza (HPAI) viruses. Recombination of HA RNA with other RNA molecules of host or virus origin is a dominant mechanism of multi basic cleavage site (MBCS) acquisition for H7 subtype HA. Using alignments of HA RNA sequences from documented cases of MBCS insertion due to recombination, we show that such recombination with host RNAs is most likely to occur at particular hotspots in ribosomal RNAs (rRNAs), transfer RNAs (tRNAs) and viral RNAs. The locations of these hotspots in highly abundant RNAs indicate that RNA recombination is facilitated by the binding of small nucleolar RNA (snoRNA) near the recombination points.

## INTRODUCTION

Low pathogenic avian influenza (LPAI) viruses, which circulate in wild bird populations, occasionally evolve into highly pathogenic avian influenza (HPAI) viruses in poultry that can cause devastating outbreaks (reviewed in Lee et al. 2020). HPAI strains emerge from LPAI viruses upon conversion of a cleavage site in the surface protein hemagglutinin (HA) from the motif recognized by trypsin-like proteases into a so-called multiple basic cleavage site (MBCS) which is recognized by furin-like proteases (Garten et al. 2015). As cleavage of the HA polyprotein precursor HA0 into two chains, HA1 and HA2, is an essential step in the virus replication cycle and furin-like proteases are present in a broader range of host tissues, the motif change facilitates the systemic virus dissemination in chickens, leading to severe disease.

Influenza A viruses are members of the *Orthomyxoviridae* family. They have a segmented negative-sense RNA genome that consists of eight segments. The viruses are classified into subtypes defined by the HA and neuraminidase (NA) proteins. Although sixteen HA subtypes (H1-H16) are known to circulate in birds, only H5 and H7 subtypes have been observed to evolve towards a HPAI phenotype in poultry (Abdelwhab et al. 2013; Lee et al. 2020). Three molecular mechanisms of the changes leading to MBCS formation have been described (reviewed in Abdelwhab et al. 2013; Naguib et al., 2019; Lee et al. 2020): (1) duplications of the nucleotides coding for the cleavage site by polymerase stuttering (common for both H5 and H7 subtypes); (2) point substitutions (usually in combination with duplications, but also observed in their absence in H5 HA segments) and (3) recombination of H7 HA RNA segment with other RNAs (Figure 1).

**Figure 1.**
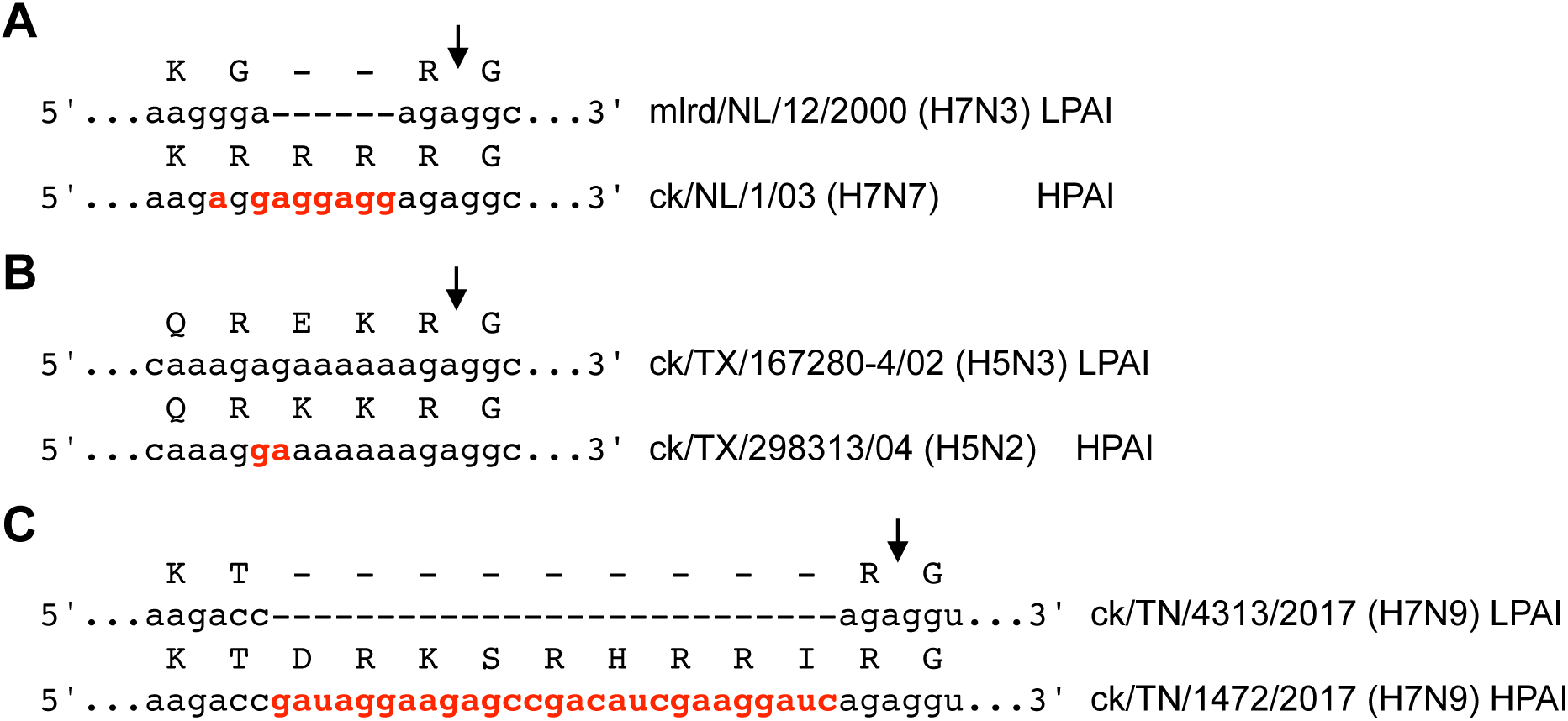
Examples of MBCS insertions in HPAI viruses. (A) Substitutions and duplications of Arg codons in the HA RNA from HPAI strain A/chicken/Netherlands/1/03 (H7N7) as compared to closely related HA RNA from LPAI virus A/mallard/Netherlands/12/2000(H7N3) (Fouchier et al. 2004). (B) Substitution creating the MBCS site of A/chicken/TX/298313/2004(H5N2) as compared to LPAI strain A/chicken/Texas/167280-4/2002(H5N3) (Lee et al. 2005). (C) Recombination of HA RNA with 28S rRNA leading to the insertion of 9 codons in the HPAI virus A/chicken/Tennessee/17-007147-2/2017(H7N9) as compared to related LPAI strain A/chicken/Tennessee/17-007431-3/2017(H7N9) (Lee et al. 2017). Inserted or mutated nucleotides are in red, locations of HA0 cleavage sites are shown by arrows.

So far, recombinations have only been observed in H7 HA segments. In a few cases with relatively large insertions (at least 21 nucleotides, or 7 codons), their origins could be reliably established by sequence database similarity searches. These insertions either originated from other segments of the virus (Orlich et al. 1994; Suarez et al. 2004; Hirst et al. 2004) or from 28S rRNA (Khatchikian et al. 1989; Maurer-Stroh et al. 2013; Lee et al. 2017) (Table 1). The origins of the shorter insertions are more difficult to identify, in particular due to the low significance of searches with short query sequences which are expected to yield false-positive hits identical to the queries. Nevertheless, some of these shorter insertions are unlikely to be produced by polymerase stuttering or sequential substitutions and rather could be due to recombination (Banks et al. 2001; Monne et al. 2014; Berhane et al. 2009; Beerens et al. 2020).

**Table 1.**
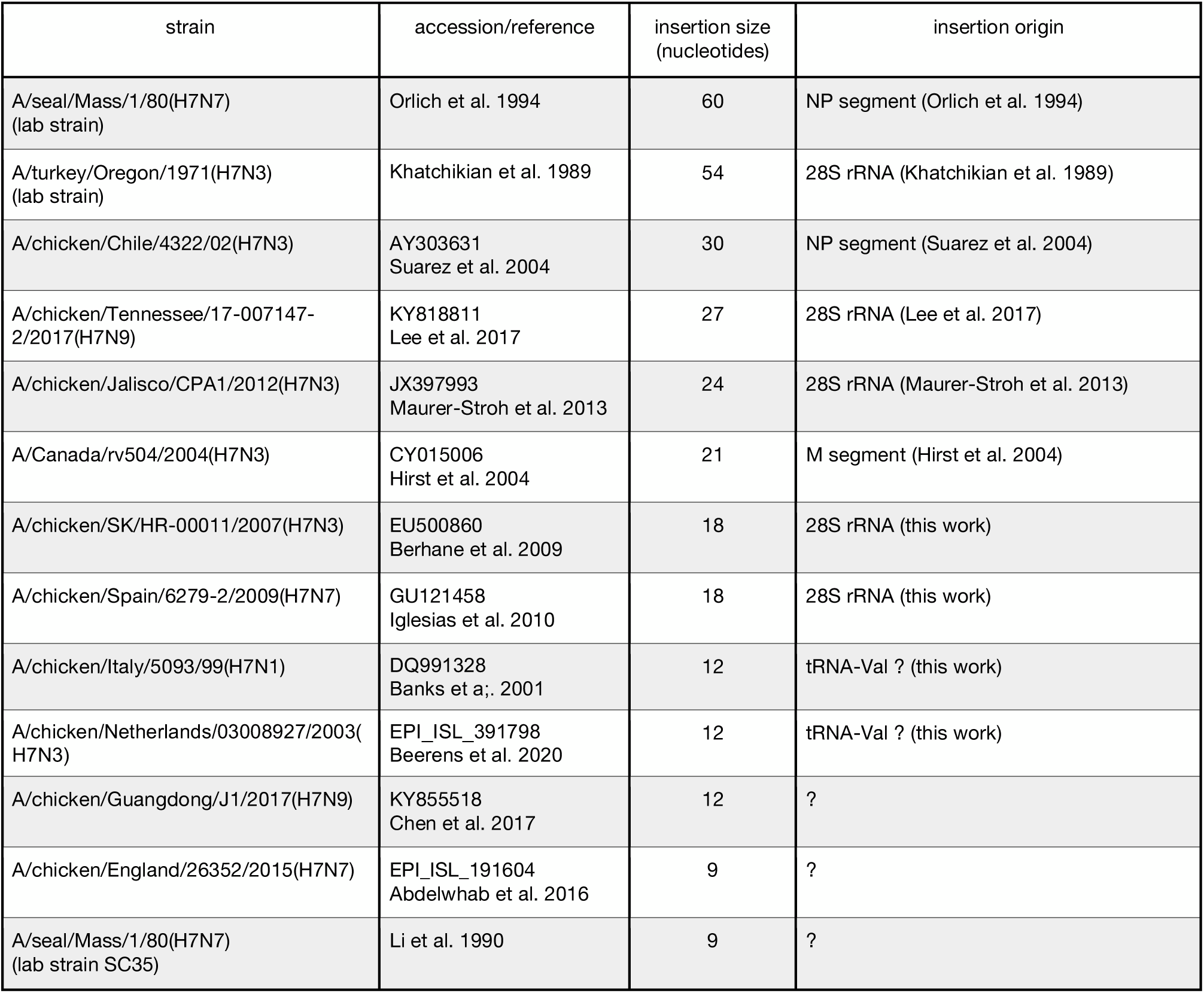
MBCS insertions into H7 HA segments with (possible) recombination origin. Representative strains of the outbreaks and viruses obtained by passaging in laboratory are listed in the descending order of insertion sizes.

Here, we used various alignment programs and database similarity searches to identify potential sources of insertions that have created MBCS sequences in known cases of H7 HPAI strains. The results led us to hypothesize that the hotspots for rRNA recombination with HA are determined by small nucleolar RNA (snoRNA) binding. Furthermore, recombination of other RNA molecules with HA also occurs at the sites homologous to snoRNA target sequences, yielding new mechanistic insight into emergence of HPAI strains from LPAI virus precursors via recombination.

The main function of snoRNAs is to guide modifications of nucleotides in ribosomal RNAs (rRNAs) and small nuclear RNAs (snRNAs) (Smith and Steitz 1997; Bachellerie et al. 2002), via base-pairing of guide regions, also called antisense elements, to target sequences. SnoRNAs are classified into two major types on the basis of their characteristic motifs and proteins found in the small nucleolar ribonucleoprotein complexes (snoRNPs): C/D box and H/ACA box snoRNAs. The C/D box snoRNPs catalyze 2’-O-methylation of nucleotides located within targets bound by antisense elements of 7-21 nucleotides. The H/ACA box snoRNPs guide pseudouridylation of uridines. Each of the guide elements of H/ACA box snoRNAs forms two relatively short duplexes. Binding of some snoRNAs to their targets is involved in regulation of processes other than nucleotide methylation (Bachellerie et al. 2002; Bratkovič et al. 2019; Bergeron et al. 2020), and snoRNA functions could be exploited by viruses (reviewed by Stamm and Lodmell, 2019). The structures of multiple snoRNAs, their guide regions and binding sites are known (Lestrade and Weber 2006; Bratkovič et al. 2019) or can be identified by sequence comparison between different species. In particular, we analyzed the binding sites of chicken snoRNAs to 28S rRNA in order to identify the features that could distinguish those located near hotspots of recombination with HA from others.

## RESULTS

The origins of insertions of 18 nucleotides in the HA segments of H7 HPAI strains isolated in Canada in 2007 (A/chicken/SK/HR-00011/2007 (H7N3)) and in Spain in 2009 (A/chicken/Spain/6279-2/2009 (H7N7)) have so far remained unidentified (Berhane et al. 2009; Iglesias et al. 2010). These insertions are the same in both HA sequences (accessions EU500860 and GU121458, respectively), although the viruses have distinct origins. Alignment of the inserted sequence with chicken rRNAs (accessions KT445934 and XR_003078040) using the Waterman-Smith algorithm showed that it was identical to the chicken 28S rRNA region 4357-4374. A BLAST search in the GenBank Nucleotide collection (nr/nt) retrieved the same identity, accompanied by multiple hits to homologous rRNA fragments in other species that are 100% conserved. The E-value of 6.5, calculated by the BLAST program for these hits, defined them as non-significant, as this value corresponds to the number of hits with at least a given score, expected by chance alone. However, a BLAST search with the same 18 nucleotides against the reference chicken genome yielded the same alignment to 28S rRNA with a considerably lower E-value of 0.027, with no other hits with 100% identity. Thus, in the reasonable assumption that the insertions resulted from HA recombination with one of the host (chicken) RNAs, one can conclude that both recombination events leading to the 2007 Canadian and 2009 Spanish HPAI involved the 28S rRNA.

Together with the previously identified cases (Khatchikian et al. 1989; Maurer-Stroh et al. 2013; Lee et al. 2017), this brings the total number of recorded rRNA recombination events of HA segments with 28S rRNA to five (Table 1). As observed in the 2007 Canadian and 2009 Spanish HPAI, the insertion observed in A/chicken/Tennessee/17-007147-2/2017 (H7N9) virus involved the same recombination site as in the Mexican H7N3 HPAI viruses (Maurer-Stroh et al. 2013; Lee et al. 2017), Therefore, two regions of rRNA were involved in four independent HPAI emergences, suggesting that recombination sites in the 28S rRNA are not random.

Interestingly, all three recombination points in the 28S rRNA turned out to be located near the known snoRNA binding sites (Figure 2A-E): either one nucleotide downstream of a region paired to a C/D box snoRNA, or just adjacent to the region paired to H/ACA box snoRNA. The probability of this to be due to chance alone is low: with 105 identified snoRNA targets (see Materials and Methods and Supplemental Table S1) in chicken 28S rRNA of 4437 nucleotides, the probability of three independent sites to be at or one nucleotide downstream of a snoRNA/rRNA duplex is (2 × 105 / 4437)^3^ ≈ 10^−4^.

**Figure 2.**
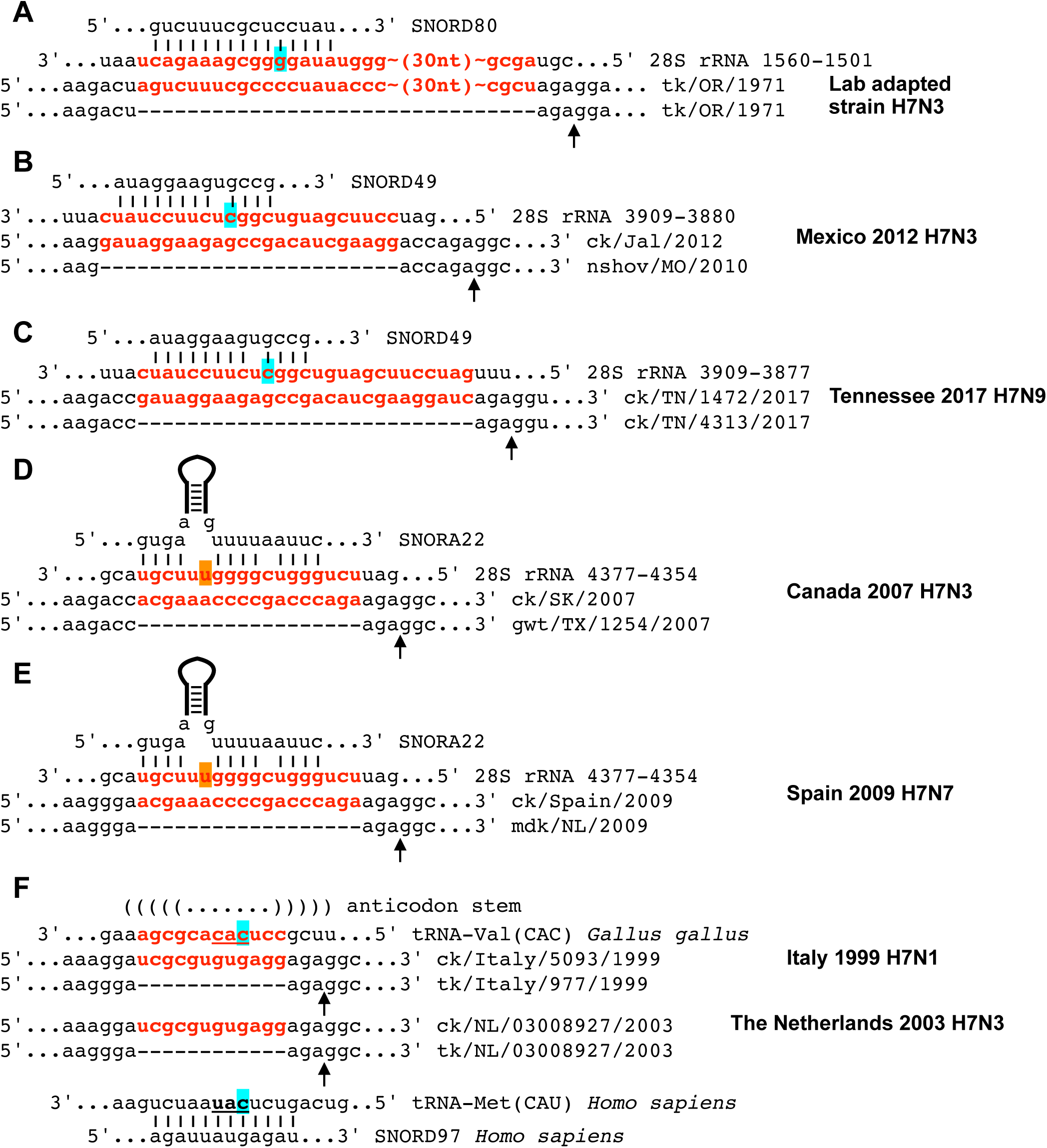
Recombination of the influenza virus H7 HA RNA with host RNAs. (A-E) The insertions in the HA segments created by recombination with 28S rRNA and the binding sites of snoRNAs near the recombination sites. The HA RNA sequences from HPAI strains (Khatchikian et al. 1989; Maurer-Stroh et al. 2013; Lee et al. 2017; Berhane et al. 2009; Iglesias et al. 2010; Banks et al. 2001; Beerens et al. 2020) are aligned to HA sequences of most closely related LPAI viruses. The apical stem between two parts of the SNORA22 antisense element is shown schematically. (F) The 12-nt sequence inserted in the Italian H7N1 (1999) and Dutch H7N3 (2003) HPAI strains is complementary to the region encompassing anticodon of chicken tRNA-Val. Base pairs of the tRNA-Val anticodon stem are shown with bracket view. Anticodon nucleotides are underlined. Chicken tRNA-Val is aligned to human tRNA-Met in order to illustrate the location of the C/D box snoRNA binding site in the tRNA-Met (Vitali and Kiss 2019). Inserted sequences and their complements in 28S rRNA and tRNA are shown in red. Locations between the codons for conserved amino acids Arg and Gly, corresponding to the HA0 polyprotein cleavage sites, are indicated by arrows. The 2’-O-methylated nucleotides are labeled with cyan, the pseudouridylated uridines with orange. For strain abbreviations, see Materials and Methods.

The optimal Waterman-Smith alignments of the insertions of 9-12 nucleotides in H7 HA segments of HPAI viruses from Italy, 1999-2001 (H7N1) (Banks et al. 2001; Monne et al. 2014), England, 2015 (H7N7) (Abdelwhab et al. 2016), China, 2017 (H7N9) (Chen et al. 2017) or laboratory-adapted H7N7 strain SC35 (Li et al. 1990) with chicken rRNAs and other segments of the corresponding strains contained mismatches and therefore did not allow one to draw conclusions on the origins of these insertions. Using the insertions of 9 nucleotides for BLAST searches in the chicken genome was not possible because such short queries were rejected by the BLAST program. It is possible to estimate the E-value of such a search in chicken genome, using a general expression E = K×m×n×exp(-λS) for gap-free alignments with the scores of at least S, where m and n are the lengths of aligned sequences and K and λ depend on scoring parameters and nucleotide composition (see e.g. Altschul et al. 1990). Thus, the pairwise BLAST alignment of the insertion of 9 nucleotides in the HA of A/chicken/England/26352/2015(H7N7) versus the full-length HA sequence of 1704 nucleotides (accession EPI_ISL_191604) yielded E-value of 0.015.

As the HA sequence is not characterized by a biased nucleotide composition, the E-value for a search in the chicken genome assembly GRCg6a (accession GCA_000002315.5) with ungapped length of 1055.6 Mb can be estimated as proportional to the genome length. The obtained E-value of more than 9000 confirmed that such BLAST search with a 9 nucleotides sequence makes no sense.

BLAST searches with the insertions of 12 nucleotides of Chinese H7N9 and Italian H7N1 strains in the reference chicken genome yielded 100%-identity hits with a reported E-value of 105. However, the actual number of hits was significantly lower than 105: 14 (taking into account two alternative variants differing by one-nucleotide shift) and 12, respectively, with only 10 corresponding to transcribed gene regions in each case. Remarkably, 6 out of 10 hits of the Italian H7N1 insertion were located in tRNA-Val(CAC) genes, suggesting that the insertion most likely originated from one of them, with the sequence complementary to the anticodon loop and part of the anticodon stem (Figure 2F). Furthermore, the same sequence was inserted into the HA RNA of LPAI virus precursors in turkeys during an outbreak in three Dutch farms in 2002 and selected rapidly upon experimental infections of chickens in 2003 (Velkers et al. 2006; Beerens et al. 2020) (Figure 2F).

One of the important factors explaining why rRNAs, and probably tRNAs are relatively frequently involved in recombination with influenza HA RNA is the high abundance of these RNAs in eukaryotic cells. Indeed, about 85-90% of the total mass of transcripts is rRNA molecules, followed by tRNAs (up to 10%), the latter being the most abundant RNA type in terms of number of molecules (e.g. Waldron and Lacroute 1975; Boivin et al. 2018). Recombination with influenza virus RNA from other genes may similarly be explained by their high abundance in virus-infected cells. Furthermore, 18S and 28S rRNAs have been shown to be packaged into influenza A virions (Noda et al. 2018) and thus have a high probability to be in close proximity to the vRNA. SnoRNAs are also highly abundant RNA species, and are frequently used by influenza A virus polymerase in cap-snatching during transcription (Koppstein et al. 2015; Gu et al. 2015; Sikora et al. 2017), although it should be noted that all three snoRNAs binding to the 28S rRNA recombination sites (Figure 2 A-E) are encoded in introns of other genes and therefore uncapped.

A mere abundance of snoRNAs cannot explain why the hotspots of rRNA recombination with the influenza A virus HA RNAs are adjacent to snoRNA binding sites. Rather, the formation of snoRNA/rRNA duplexes seems to play an important role in the recombination mechanism. The fact that two out of more than hundred snoRNA binding sites in 28S rRNA were repeatedly involved in four independent recombination events leading to HPAI phenotype (Fig.2, B-E) might be determined by (1) favorable properties of oligopeptides encoded by these regions upon insertion into the hemagglutinin protein, (2) relative abundance of these snoRNAs as compared to others, (3) certain features of snoRNA/rRNA duplexes that mediate recombination and/or (4) effects of HA RNA structures that are formed upon insertions on virus replication. These points were addressed one-by-one below.

We did not identify any clear pattern in oligopeptides inserted into the HA sequences upon recombination that could explain the preference for locations near binding sites of SNORD49 and SNORA22 (Figure 3). Of course, these amino acid sequences matched the requirements for furin cleavage, but so did a number of oligopeptides that could be inserted if recombination would occur at similar locations near the binding sites of other snoRNAs (Supplemental Table S1). Using the main furin cleavage site requirements for arginine at position P1 and at least two basic residues at positions P2, P4 and P6 (Duckert et al. 2004), we identified 69 such amino acid sequences with lengths up to 20 residues (Supplemental Table S1), which is the maximal length of documented insertions (Table 1), encoded by regions at the binding sites of 29 snoRNAs. According to the estimates of furin cleavage efficiencies using a neural network - based algorithm (Duckert et al. 2004), many of these sequences would be cleaved even more efficiently than those inserted in the natural HPAI strains (Figure 2). The amino acid sequences encoded by the 28S rRNA regions at the SNORD49 and SNORA22 binding sites obtained scores of 0.625 and 0.658, respectively, while higher scores were calculated for e.g. the sequences encoded by the regions of SNORD35 and SNORD104 binding: 0.779 and 0.685, respectively. As such scores are used to quantify the cleavage efficiency (Duckert et al. 2004), one has to conclude that the HA insertions of rRNA regions at the SNORD49 and SNORA22 binding sites did not create the most efficiently cleaved sequences. Apart from furin cleavage requirements, no other motif was found in the amino acid sequences inserted in the HA protein by recombination (Figure 3). No difference in potential distortion of HA protein three-dimensional structure was detected either: homology modeling of the protein structures with both naturally observed rRNA insertions and those derived from other sites (not shown) using the SWISS-MODEL algorithm (Waterhouse et al. 2018) predicted no considerable conformational changes in the regions flanking the cleavage site, which is located in the exposed loop (Chen et al. 1998). This was consistent with the fact that diverse inserted sequences can be accommodated at the H7 HA cleavage site upon recombination (Abdelwhab et al. 2013). Of course, we cannot exclude that certain combinations of HA cleavage loop sizes and sequences could be more favorable for virus replication.

**Figure 3.**
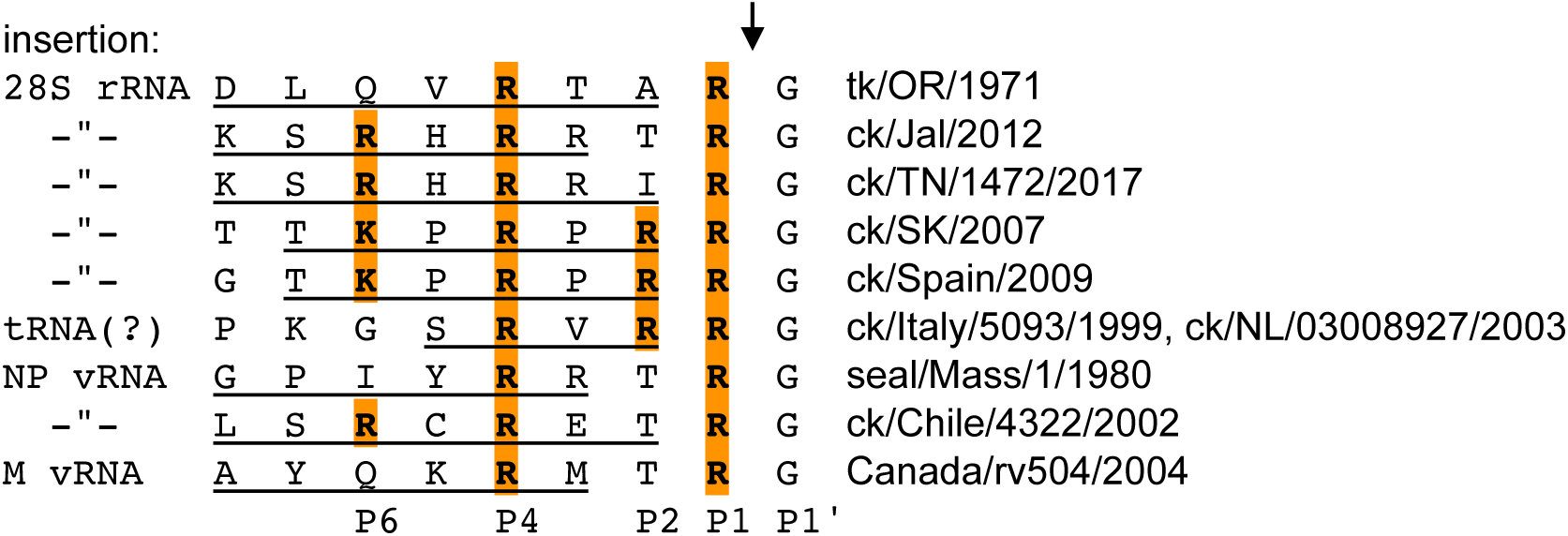
MBCS amino acid sequences in H7 HA RNAs yielded by recombination (Khatchikian et al. 1989; Maurer-Stroh et al. 2013; Lee et al. 2017; Berhane et al. 2009; Iglesias et al. 2010; Banks et al. 2001; Beerens et al. 2020; Orlich et al. 1994; Suarez et al., 2004; Hirst et al. 2004). Amino acids at eight positions upstream of the cleavage site (P1-P8) which could affect cleavage by furin-like proteases (Duckert et al. 2004) are shown, inserted amino acids are underlined. Important basic residues at positions P1, P2, P4 and P6 are highlighted, cleavage site is shown by arrow. For strain abbreviations, see Materials and Methods.

The three chicken snoRNAs suggested here to be involved in rRNA recombination with HA segments (Figure 2) are not among the most abundant ones according to expression levels reported in the Expression Atlas (www.ebi.ac.uk/gxa/home). Among snoRNAs potentially leading to furin MBCS insertions following the model discussed above, they rank between 3 and 21 out of 21 snoRNAs for which expression data have been generated (Merkin et al. 2012; Barbosa-Morais et al. 2012), depending on the tissue and the study (see Supplemental Table S1). The best rank (3) was obtained for SNORD49 in skeletal muscle tissue according to Merkin et al. (2012), though according to the data of Barbosa-Morais et al. (2012) this rank is 5. The ranks of SNORA22 in this tissue were 21 and 13 in the two studies. It should be noted, however, that snoRNA levels can change upon infection by viruses, in particular, influenza viruses (Peng et al. 2011; Stamm and Lodmell 2019; Samir et al. 2019).

Thermodynamic stability is one of the features of snoRNA/rRNA duplexes that could affect their function. However, no general trend was observed in the free energies of the binding of the three snoRNAs suggested here to trigger 28S rRNA recombination with HA RNA (Figure 2, Supplemental Table S1). The binding of SNORD80 to its site was one of the strongest with −28.1 kcal/mol, which was the fourth strongest duplex out of 53 chicken C/D box snoRNA duplexes formed with 28S rRNA, ranging between −28.8 and −12.0 kcal/mol. However, the rank of SNORD49 pairing to rRNA was only 29 with −22.4 kcal/mol, and SNORA22 formed the weakest binding duplexes out of 52 identified binding sites of H/ACA box snoRNAs to 28S rRNA: −3.8 and −3.0 kcal/mol.

The presence of mismatches was a feature of snoRNA/rRNA duplexes that seemed to correlate with repeated involvement of the 28S rRNA sites near SNORD49 and SNORA22 binding in the recombination with HA RNA (Figure 2). Only five other chicken snoRNAs, namely, SNORD12/SNORD106, SNORD24, SNORD50, SNORA17 (GGN58) and SNORA63, formed duplexes with mismatches upon binding to 28S rRNA. Moreover, out of these five duplexes only the SNORA17 pairing and subsequent recombination with HA RNA would result in a furin cleavage signal in the HA polyprotein. A mismatch in the snoRNA/rRNA pairing interaction might play a role in rRNA recombination at the binding site due to a faster dissociation of the snoRNA, which would be required to use the binding region as a template by the polymerase. Kinetics of snoRNA release from the rRNA recombination site can be important to allow for template switching during a time window of influenza polymerase slowdown, which is likely to occur in the cleavage site region due to its structured character (Gultyaev et al. 2019).

Structural constraints in this HA RNA domain (Gultyaev et al. 2016; 2019) may also determine the fitness of the viruses containing the insertions. The insertions occur in the proposed hairpin loop of the conserved H7 HA RNA structure, stabilizing it by base pairs formed by inserted sequences (Gultyaev et al. 2016). However, predictions of possible structures of this domain with inserted sequences from other rRNA regions showed similar stabilization (not shown), so that we could not derive any specific feature characteristic for SNORD49 and SNORA22 targets, which would explain their repeated involvement in the recombination with HA RNA.

Interestingly, we identified chicken snoRNAs that could bind to the known sites (Table 1) of recombination of NP and M segments with HA vRNA (Figure 4). As a number of snoRNA functions are determined by their targeting various RNAs (Bachellerie et al. 2002; Bratkovič et al. 2019; Bergeron et al. 2020), it is possible to suggest that emergence of HPAI strains by HA recombination with RNA molecules other than rRNA can be also triggered by snoRNA binding.

**Figure 4.**
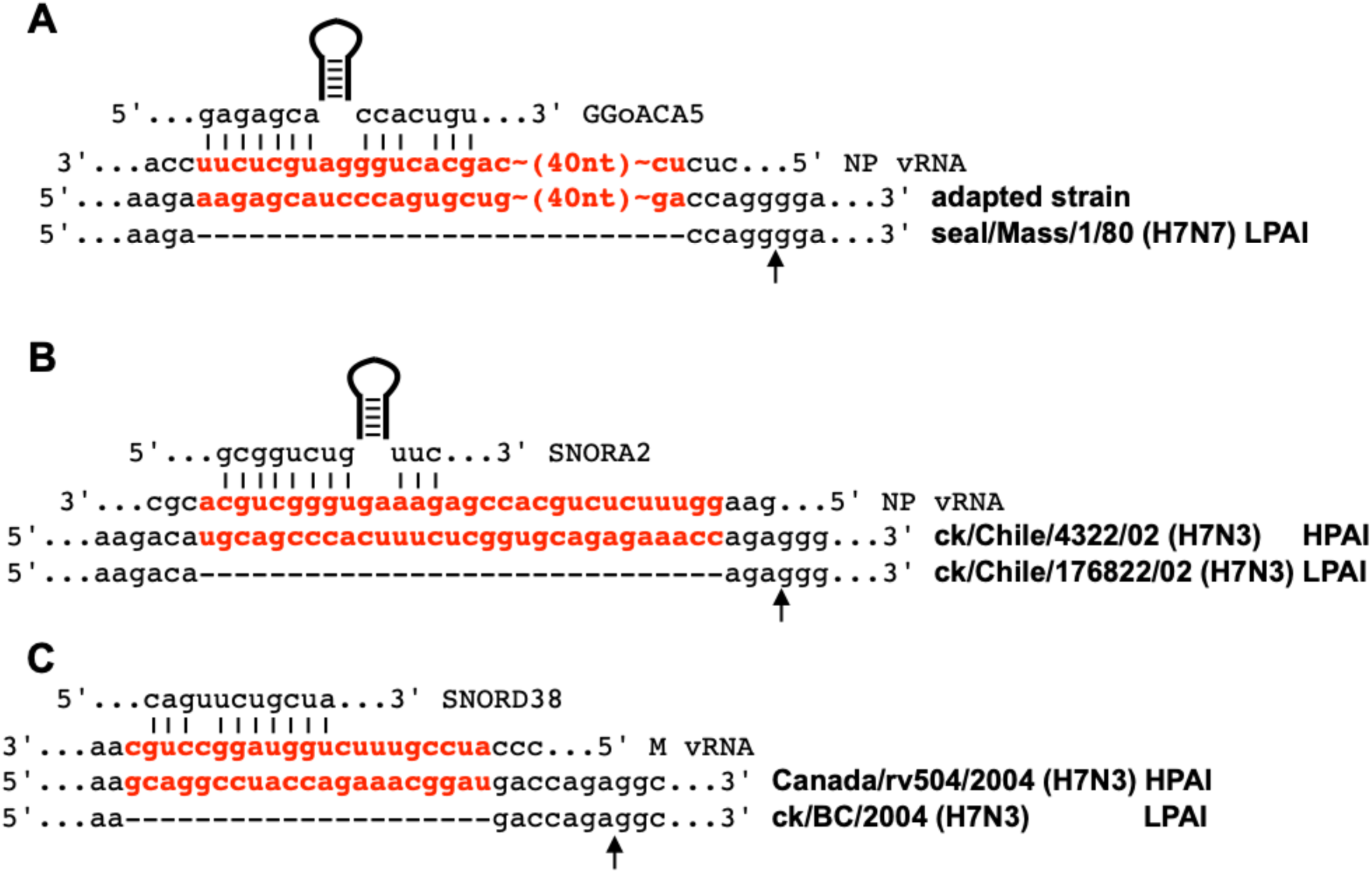
Putative snoRNA binding at the recombination sites of HA RNAs with other influenza A virus segments. The sequences of NP and M segments involved in the recombination are from the LPAI progenitors (Orlich et al. 1994; Suarez et al., 2004; Hirst et al. 2004). Chicken orphan H/ACA snoRNA GGoACA5 has been identified by Shao et al. (2009). Notations are similar to those in Figure 2.

## DISCUSSION

Here, we identified that for all five described cases of recombination of H7 HA RNA with chicken rRNA, the recombination point was located near snoRNA binding sites. Interestingly, two snoRNAs (SNORD22 and SNORD49) were involved in two independent cases of recombination. We therefore hypothesize that recombination of HA segments with rRNA occurs at the hotspots determined by snoRNA/rRNA duplexes. Recombination may be facilitated by polymerase slowdown induced by the template secondary structure around the recombination site (Gultyaev et al. 2019). Upon release of snoRNA, the rRNA is exploited as a template by a copy-choice mechanism, starting from the 3’ end of the snoRNA binding site. After the second template switching back to the viral template RNA, a nascent viral cRNA with an rRNA-derived insertion is produced. If the insertion encodes a MBCS at the HA1/HA2 boundary in the encoded polypeptide and does not considerably interfere with virus fitness, a highly pathogenic strain can emerge.Presumably, similar recombination events may occur in other regions of HA RNA, but they remain undetected because distorted HA protein structures make the resulting virions not viable.

Mechanistic details of such a recombination remain unclear. During influenza virus RNA replication the template and nascent strand regions near the elongation point are located in the channels formed by the polymerase subunits (te Velthuis and Fodor 2016; Pflug et al. 2017; Fodor and te Velthuis, 2019). Recombination would probably require a temporary disruption of some interactions between and/or within the subunits to accommodate another template in a copy-choice mechanism. Alternatively, an involvement of another polymerase molecule cannot be excluded. It should be noted that non-canonical copying of RNA templates by DNA-dependent RNA polymerases, including rRNA-transcribing RNA polymerase I, has been observed (Macnaughton et al. 2002; Jain et al. 2020). It is also possible that the recombination hotspots at the snoRNA binding sites in rRNA and vRNA are determined by a duplex-specific cleavage that could yield the free 3’ ends of the targeted regions.

Similar to HA recombination with 28S rRNA, the suggested recombinations with tRNA in the Italian H7N1 and Dutch H7N3 HPAI strains would occur during cRNA synthesis, as in all these cases the fragments inserted into the coding sequences are complementary to host non-coding RNAs (Figure 2). Although tRNA modifications in eukaryotes are mostly snoRNA-independent, potential snoRNA targets have been identified in nematode tRNAs (Zemann et al. 2006) and a precedent of a C/D box snoRNA-tRNA interaction guiding 2’-O-methylation of wobble nucleotide in the anticodon of human tRNA has been recently set (Vitali and Kiss 2019). It is tempting to speculate that the insertions of the 12-nucleotide tRNA fragment in HA RNAs could be triggered by an interaction of the similar region in avian tRNA-Val with yet unknown C/D box snoRNA. A putative homologous binding site in chicken tRNA-Val would be located one nucleotide upstream of the terminal nucleotide of the sequence inserted in HPAI strains, exactly like the SNORD80 and SNORD49 binding sites in 28S rRNA (Figure 2 A, B, C, F). Although we did not identify any chicken snoRNA with a guide sequence that could target the tRNA-Val anticodon, such a snoRNA may exist as many snoRNAs remain unannotated (Gardner et al. 2010; de Araujo Oliveira et al. 2016; Bratkovič et al. 2019; Bergeron et al. 2020)

If true, the preference for snoRNA-bound locations in template switching by the influenza A virus polymerase is an important part of the mechanism of the emergence of HPAI H7 strains by recombination of virus HA segments with other RNAs. The hypothesis provides a rationale behind the existence of hotspots in recombination with rRNA. Future experimental studies could elucidate the details of this mechanism.

## MATERIALS AND METHODS

### Sequences and alignments

The strains of HPAI viruses yielded by HA recombination and their likely LPAI progenitors were identified from literature (Khatchikian et al. 1989; Maurer-Stroh et al. 2013; Lee et al. 2017; Berhane et al. 2009; Iglesias et al. 2010; Banks et al. 2001; Monne et al. 2014; Beerens et al. 2020; Orlich et al. 1994; Suarez et al., 2004; Hirst et al. 2004) and/or BLAST (Camacho et al. 2009) searches. Accessions: A/turkey/Oregon/1971(H7N3) (abbreviated here as tk/OR/1971), M31689; A/chicken/Jalisco/CPA1/2012(H7N3) (ck/Jal/2012), JX397993; A/northern/shoveler/Missouri/10OS4750/2010(H7N3) (nshov/MO2010), CY133416; A/chicken/Tennessee/17-007147-2/2017(H7N9) (ck/TN/1472/2017), KY818811; A/chicken/Tennessee/17-007431-3/2017(H7N9) (ck/TN/4313/2017), KY818819; A/chicken/SK/HR-00011/2007(H7N3) (ck/SK/2007), EU500860; A/green-winged teal/TX/1254/2007(H7N3) (gwt/TX/1254/2007), KF573723; A/chicken/Spain/6279-2/2009(H7N7) (ck/Spain/2009), GU121458; A/mallard duck/Netherlands/2/2009(H7N7) (mdk/NL/2009), KX978114; A/chicken/Italy/5093/99(H7N1) (ck/Italy/5093/1999), DQ991328; A/turkey/Italy/977/1999(H7N1) (tk/Italy/977/1999), GU052999; A/chicken/Netherlands/03008927/2003(H7N3) (ck/NL/03008927), EPI_ISL_391798; A/turkey/Netherlands/03008927/2003(H7N3) (tk/NL/03008927), EPI_ISL_391799; A/chicken/Chile/4322/02(H7N3) (ck/Chile/4322/02), AY303631; A/chicken/Chile/176822/02(H7N3) (ck/Chile/176822/02), AY303630; A/Canada/rv504/2004(H7N3) (Canada/rv504/2004), CY015006; A/GSC_chicken/British Columbia/04(H7N3) (ck/BC/2004), AY650270. Chicken 28S rRNA sequence: accession XR_003078040. Sequences of tRNAs were retrieved from the tRNAdb database (Jühling et al. 2009). Alignments of insertions versus rRNA or influenza virus RNAs were carried out by the Smith-Waterman algorithm of the program Water provided by the EMBL-EBI tools (Madeira et al. 2019).

Available chicken snoRNAs that target 28S rRNA were initially retrieved from the snoRNA databases: snoRNABase (Lestrade and Weber 2006) and snOPY (Yoshihama et al. 2013). This list was complemented by published records of chicken snoRNA sequences (Shao et al. 2009; Zhang et al. 2009; Deryusheva et al. 2020). The final non-redundant dataset of chicken snoRNA sequences and structures was verified by identification of GenBank database entries, genomic sequences (using BLAST searches in the chicken genome) and structural models from the database of RNA families Rfam (Kalvari et al. 2018). In case of alternative names for the same snoRNA, the symbols approved by the HUGO Gene Nomenclature Committee (HGNC) (Braschi et al. 2019) were preferably used, that is, SNORD… and SNORA… for C/D and H/ACA box snoRNAs, respectively. Sequence accessions of the snoRNAs that bind to 28S rRNA near the points of recombination with HA RNA: SNORD80 (or snR60), XR_003076593; SNORD49 (or GGgCD65), XR_003071336; SNORA22 (or GGN50), XR_003071339. SnoRNAs that may bind to NP and M segments near the points of recombination: SNORD38, XR_003076587; SNORA2, NC_008465 (chromosome 33), positions 3350951-3351091 (complement); GGoACA5, NC_006112 (chromosome 25), 3660678-3660803 (complement).

### Analysis of snoRNA-rRNA duplexes

The binding sites of chicken snoRNAs in 28S rRNA were identified by analogy to duplexes identified for human snoRNAs, given in the snoRNABase (Lestrade and Weber 2006) and using the identification of guide sequences in snoRNAs that are complementary to chicken 28S rRNA. Free energies of RNA duplexes were calculated using thermodynamic parameters from the Nearest Neighbor Database (Turner and Mathews 2010).

### SnoRNA expression levels

The expression levels of chicken snoRNAs were obtained from the data given by Expression Atlas incorporated in the Ensembl genomic browser (Papatheodorou et al. 2018; Yates et al. 2020).

## ACKNOWLEDGEMENTS

This work was supported by NIAID/NIH contract HHSN272201400008C. We gratefully acknowledge the originating and submitting laboratories for sequences in the GISAID’s EpiFlu Database, used in this study.

